# Prognostic Role of c-Met Overexpression in High Grade Glioma: a Meta-analysis

**DOI:** 10.1101/695775

**Authors:** Bo Wu, Yuhan Ma, Sheng Zhong, Junliang Ge, Shanshan Jiang, Yuan Zhang, Haiyang Xu

## Abstract

**Objectives:** This study aims to assess the relationship between the expression of c-Met and the prognosis of high grade glioma patients.

**Method:** The MET proto-oncogene encoded c-Met protein. The gene expression data of 325 patients were downloaded from CGGA. The Oncomine database analysis and the prognosis analysis were conducted. Besides, meta-analysis was also performed to confirm the conclusion.

**Result:** Oncomine database was identified and analyzed and results showed that the MET copy number was obviously higher in glioblastoma than normal tissue consistently (p<0.001). The prognostic analysis of 325 high grade glioma samples showed that high c-Met expression patients had poor overall survival (OS) and progression free survival (PFS) than the low c-Met expression patients dramatically (HR, 2.223; 95% CI: 1.662 to 2.974; P<0.0001 and HR, 2.089; 95% CI: 1.578 to 2.770; P<0.0001). 6 studies involving 503 patients were included in the meta-analysis. The pooled results indicated that the high expression of c-Met was not significantly associated with OS (HR =1.01, 95% CI:0.93-1.09), but strongly connected with shorter PFS (HR =1.92, 95% CI:1.42-2.58, p<0.01).

**Conclusion:** c-Met overexpression has correlation with poor prognosis of high grade glioma patients.

## Introduction

Glioma is the most common primary central nervous system tumors. Based on WHO classification, it is categorized into four grades. High grade glioma, also known as malignant glioma, included grade III tumors (anaplastic astrocytoma, oligoastrocytoma and oligodendroglioma) and grade IV tumors (glioblastoma) ^1^. Glioblastoma (GBM), WHO grade IV gliomas, is the third CNS histological type which is most frequently reported. It is the most malignant and lethal histological type, with annual average age-adjusted incidence being 3.2 per 100,000 population in the United States ^2^. It has devastating pathology characteristics and highly invasive biological behavior. Currently, there is no enough efficient therapeutic options to prevent glioblastoma progression. People attempt to use many therapeutic ways to prolong patients’ lives, but without much effect. The mainstream of glioblastoma treatment is surgery and adjuvant radiotherapy combined with chemotherapy ^3^. However, there is still only 5.5% of Five-Year Relative Survival Rates ^2^. People are committed to looking for more effective ways to early identify and treat this disease. However, glioblastoma has diverse molecular characteristics, which has increase the difficulty in elucidating underlying mechanism. Although many molecular characterizations are recognized to be relevant to glioblastoma progression possibly, including PTEN, EGFR, HGF/MET signaling pathway, etc. Great efforts have been paid to gain insight into the genetic develop of glioma, the exact mechanism was still not fully understood. There is no targeted drug have been applied in clinic to prolong patients’ survival successfully ^4^.

The MET proto-oncogene is located on chromosome 7q21–31, encoding c-Met protein. MET gene amplification is associated with poor outcome in many disease ^5, 6^. Proto-Oncogene Proteins c-Met is a sort of cell surface protein-tyrosine kinase receptors for its only known ligand HGF. Hepatocyte growth factor (HGF), also called scatter sactor (SF) or hepatopoietin, is a multifunctional cytokine which plays an important role in many cell life activities. This protein is constituted by two chains, consisting of six domains: N-terminal domain, four kringle domains and a serine proteinase homology (SPH) domain ^7, 8^; As HGF receptor, c-Met is a dimer including a 50 kD extracellular α-subunit linked to 140 kD transmembrane β-subunit by a disulfide bond. The extracellular portion consists of the Sema domain, the PSI domain and IPT domains. The intracellular portion consists of juxtamembrane sequence, catalytic domain and C-terminal region. c-Met is expressed on the surface of epithelial and endothelial cells ^9, 10^. When binding to HGF and then active HGF/MET signaling pathway, they can mediate multiple downstream signal transduction pathways, including PI3K–AKT signaling pathway, MAPK signaling pathways, STAT signaling pathway, etc. So HGF/MET signaling pathway is involved in many biological processes ^9^. Tumor cells demonstrate HGF/MET dysregulation ^11^. It has been proved that inappropriate activation of the HGF-MET pathway is observed in in several different tumor types, including colorectal cancer, cervical cancer, hepatocellular carcinoma and so on. In these tumors, the dysregulation of HGF/MET always indicates aggressive cancer phenotype and poor survival ^12-14^.

Amount of previous studies have focused on the correlation between c-Met overexpression and many disease, like non-small cell lung cancer, kidney cancer, high grade glioma, patients’ prognosis ^15^. However, there is still a debate whether overexpression of c-Met indeed has correlation with poor survival of glioma patients. Some studies support this opinion ^16^, whereas other studies have given the evidence to the contrary to oppose to this view ^17^. Hence, in order to confirm the relationship between c-Met overexpression and high grade glioma patients’ prognosis, we performed this diversification analysis to solve this problem. At the DNA level, Oncomine database analysis is performed. At the mRNA level, CGGA glioma transcriptome dataset is also analyzed. At the protein level, meta-analysis is conducted to strengthen the credibility of conclusion. The framework of this study was shown in **Figure 1**.

**Figure 1.**
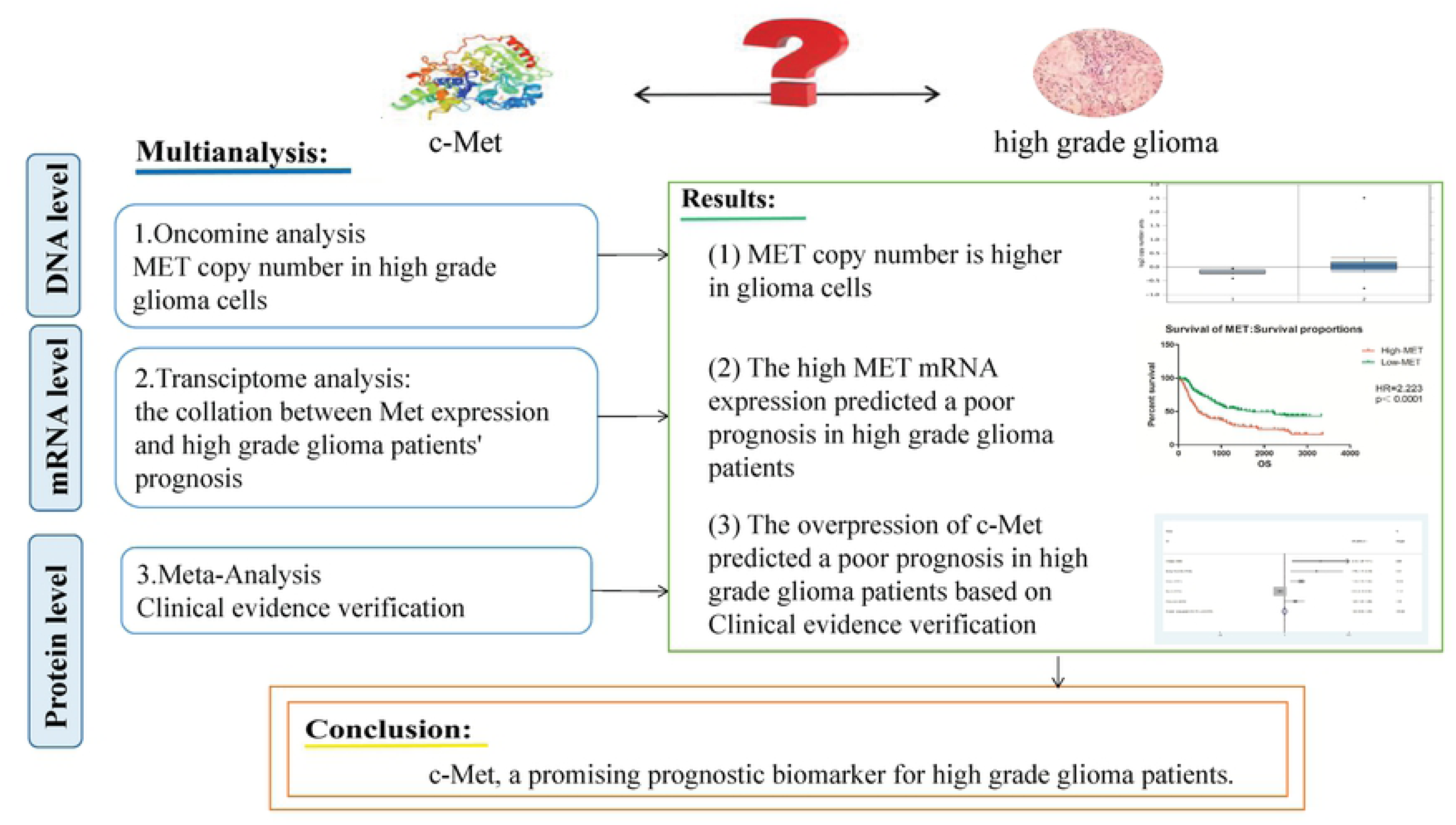
The framework of this study.

## Material and methods

### Oncomine database analysis

The Oncomine platform (http://www.Oncomine.org) is an online microarray database, from web applications to translational bioinformatics services, was applied to examine the genes’ DNA copy number difference of tumor and normal tissue in human cancers ^7, 18^. The thresholds were limited as follows: *p* value: 0.01; fold change: 2; gene rank: 10%; data type: DNA. For MET, we carried out comparisons by Glioblastoma tissue vs. Normal tissue analysis.

### CGGA glioma transcriptome dataset

The gene expression data sources, downloaded from CGGA (Chinese Glioma Genome Atlas, http://www.cgga.org.cn), included 325 patients (203 males and 122 females), with a 43.4 average age. All 325 patients enrolled were diagnosed as glioma by pathology or radiological examination. Patients’ basic characteristics including histological subtypes, WHO grade classification, age, outcomes (OS and PFS information) were all recorded. The MET proto-oncogene encoded c-Met protein. According to the MET expression level, the whole patients were split into two groups: high-MET expression group and low-MET expression group. The Kaplan-Meier analysis was performed to test whether MET expression level affected the glioma patients’ progress free survival (PFS) and overall survival (OS).

### Publication searching

The process of study selection was showed in the **Figure 2**. The relative studies were searched in PubMed, Web of Science, Embase databases up to August 20, 2018 without any language restrictions and year of publication. This searching process was executed and then cross-checked by two reviewers (Wu Bo and Ma Yuhan) independently. The searching keywords were: “Proto-Oncogene Proteins c-Met”, “MET Receptor Tyrosine Kinase”, “glioma”, “glial cell tumor”, “astrocytoma”, “Glioblastoma”, etc. The search strategy was affiliated in **Appendix 1**.

**Figure 2.**
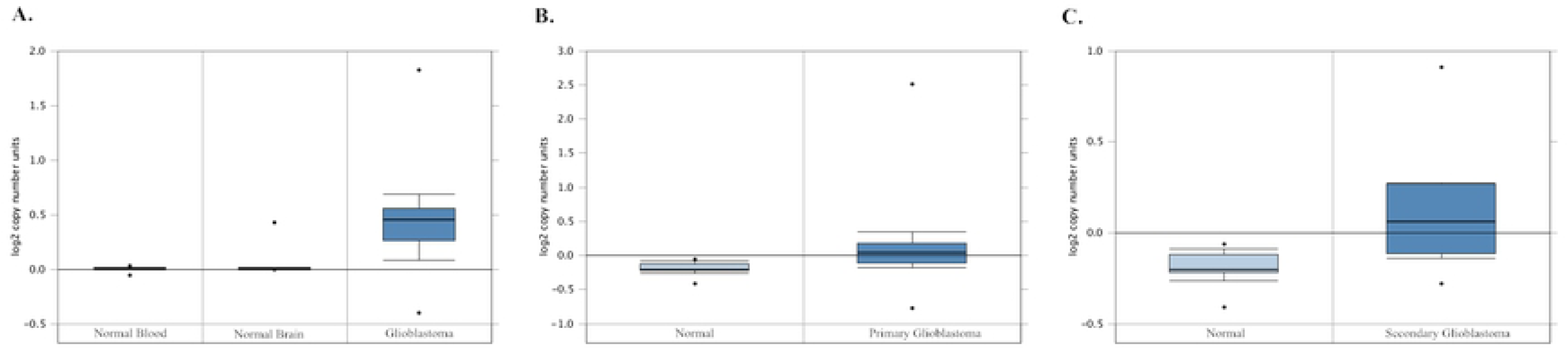
MET copy number was investigated based on Beroukhim Brain and TCGA Brain study (P<0.001). (**A**) TCGA Brain: Normal blood vs Normal brain vs Glioblastoma (cases=1531, fold change=1.338, P<0.001) (**B**) Beroukhim Brain: Normal vs Primary Glioblastoma (cases=187, fold change=1.199, P<0.001) (**C**) Beroukhim Brain: Normal vs Secondary Glioblastoma (cases=187, fold change=1.214, P<0.001)

### Study selection

Studies were considered to be eligible if they met the following criteria: (1) It has been published as original articles. (2) Articles were published as a full paper in English. (3) It is not case reports, reviews, letters or the studies with patients less than 10. (4) All study population was pathologically confirmed as having glioma by radiology or histology. (5) The article had complete information of clinical features and tumor stage information. (6) This article must investigate about the correlation between the expression of c-Met and gliomas patients’ survival. (7) There was available data or Kaplan-Meier curves about the OS or PFS of glioma patients and the control, hazard ratio (HR), 95% CI, and P value could be calculated. (8) It must be high quality papers.

According to inclusion criteria, all studies were assessed by screening the titles and abstract firstly and then read the full content of selected articles by two reviewers (Wu Bo and Ma Yuhan).

### Data extraction

The data was extracted by two reviewers (Wu Bo and Ma Yuhan) independently. General characteristics including first author, publication year, sample size, patients age, sex, method and follow-up were extracted. OS, PFS levels with HRs and corresponding CI of each study was calculated from the available data or estimated by Kaplan-Meier survival curves with the methods reported by Tierney et al ^19^. All data were checked by another author (Wo Bo and Ma Yuhan) again.

### Assessment of studies quality

The quality of all enrolled studies was assessed by using Newcastle-Ottawa Quality Assessment Scale (NOS) by two authors (Wo Bo and Ma Yuhan) independently. NOS is composed of three parts: subject selection (0-4 points), comparability of subject (0-2 points), and outcome (0-3 points). The total points of study with more than 6 scores were considered to be high quality studies. The assessment scores were showed in **Appendix 2**.

### Outcome definition

The data from each included study were recorded by two investigators (Wo Bo and Ma Yuhan) independently. The main outcome was defined as: OS (Overall Survival) was the time beginning from the date that patients had been diagnosed as glioma, to the date of patients’ death or the last follow-up visit for surviving cases. PFS (Progression Free Survival) was the time beginning from the date that patients had been diagnosed as glioma, to the date of glioma progression, death without progression, or the last follow-up visit for surviving patients without progression.

### Statistical analysis

About OS and the PFS, we used Kaplan-Meier method to estimate the differences between the two groups compared with a two-sided log-rank test in Prism GraphPad (http://www.graphpad.com/). We used STATA12.0 to calculate pooled Hazard Ratio (HR). We constructed forest plots and calculate HR with their 95% confidence interval (95% CI) to evaluate the association between c-Met expression and glioma prognosis. Data were combined according to the fixed-effect model when the heterogeneity was present. We defined that P<0.05 indicated statistical significance. I^2^ metric was used to assess statistical heterogeneity. When I^2^>50%, it would be considered significant heterogeneity. Publication bias was evaluated using the test of Begg and the test of Egger. When P-value>0.05, it would be considered as no publication bias.

## Results

### Oncomine database analysis

Oncomine database analysis was conducted to compare the DNA copy number of MET in glioma and normal samples. With the thresholds (fold change: 2; p value: 0.01; gene rank: 10%; data type: DNA), two datasets were identified and analyzed (details showed in **Table 2**). Compared with normal samples, MET copy number in glioblastoma was higher in TCGA distinctly (cases=1531, fold change=1.338, P<0.001), as well as in study Beroukhim Brain (cases=187, fold change=1.199, P<0.001) (cases=187, fold change=1.214, P<0.001) (**Figure 2**).

### Prognostic analysis on transcriptome

As shown in **Figure 3A**, the results showed that patients in the high-MET expressed group had significantly shorter OS than low-MET expressed group (HR, 2.223; 95% CI: 1.662 to 2.974; P<0.0001). Also, as **Figure 3B**, PFS was significantly shorter in the high-MET expressed group (HR, 2.089; 95% CI: 1.578 to 2.770; P<0.0001).

**Figure 3.**
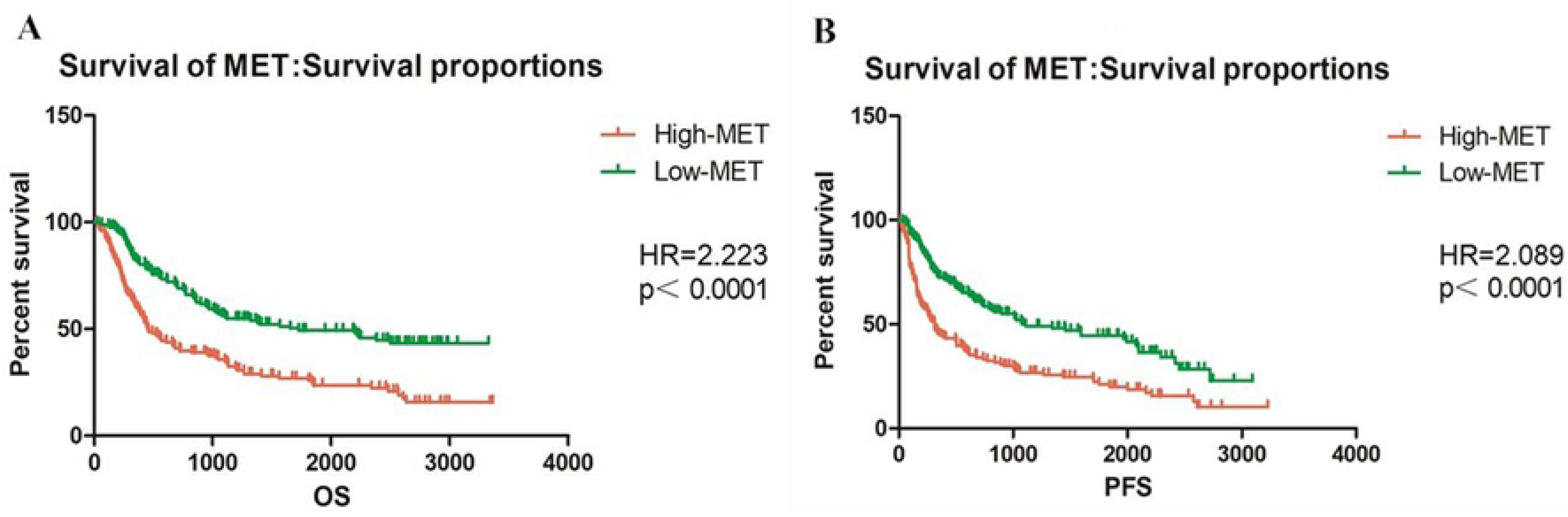
(**A**) the patients in the high-MET expressed group had significantly shorter OS than low-MET expressed group (HR=2.223; 95% CI: 1.662 to 2.974; P<0.0001). (**B**) PFS was significantly shorter in the high-MET expressed group (HR=2.089; 95% CI: 1.578 to 2.770; P<0.0001).

### Publication searching results

Through database search, we have initially identified 1837 articles and then 490 duplicates were excluded. After carefully reading, a total of 6 studies were included in the final meta-analysis. The process of selecting studies was presented in **Figure 4**.

**Figure 4.**
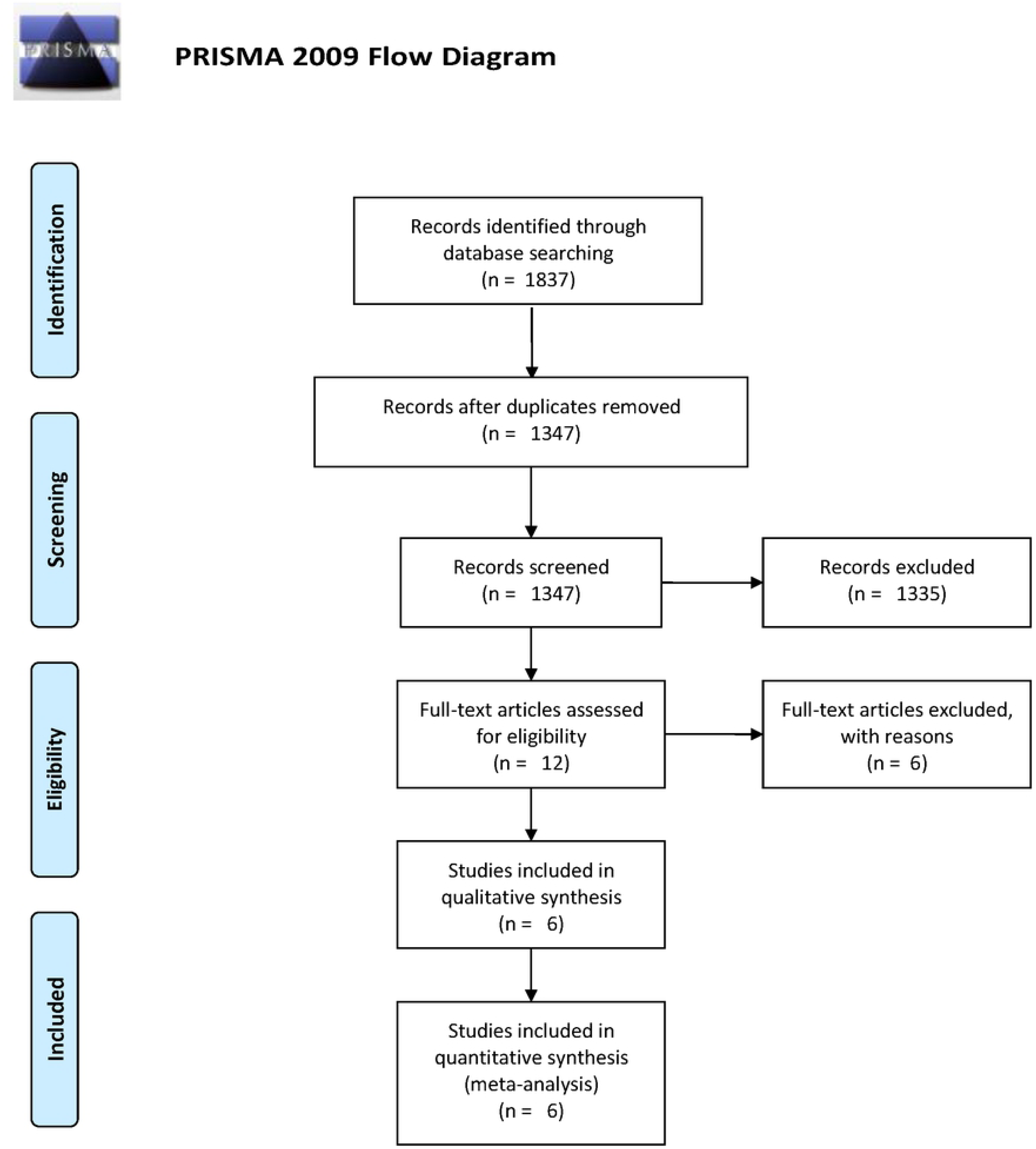
Flowchart of the study selection.

### Study characteristics and study methodological quality

Six studies ^16, 17, 20-23^, conducted in Denmark, Japan, China, Korea, Turkey, involving 503 patients were available for this meta-analysis. The published years of studies ranged from 1998 to 2015. Given the available clinical evidence, 5 articles mentioned the relationship between c-Met expression and OS of glioma patients. 3 articles described whether c-Met expression influence PFS of glioma patients. The quality of enrolled studies was assessed by Newcastle-Ottawa Scale, and the detail of scores was shown in **Appendix 2**, which showed these studies had high quality. The characteristics of the included studies were listed in **Table1**.

### Meta-analysis results

#### Overall survival

Five studies reported OS data of the relationship between c-Met expression and survival of glioma patients. In this meta-analysis, the pooled results indicated that the overexpression of c-Met was not associated with overall survival (HR =1.01, 95% CI:0.93-1.09). Significant heterogeneity was observed for the high I^2^ values (I^2^=91.3%). **(Figure 5)** Considering the HR and weight of each study, Kwak 2015 thoroughly deviated from the other studies in researches, which indicated that it may be the cause of the heterogeneity.

**Figure 5.**
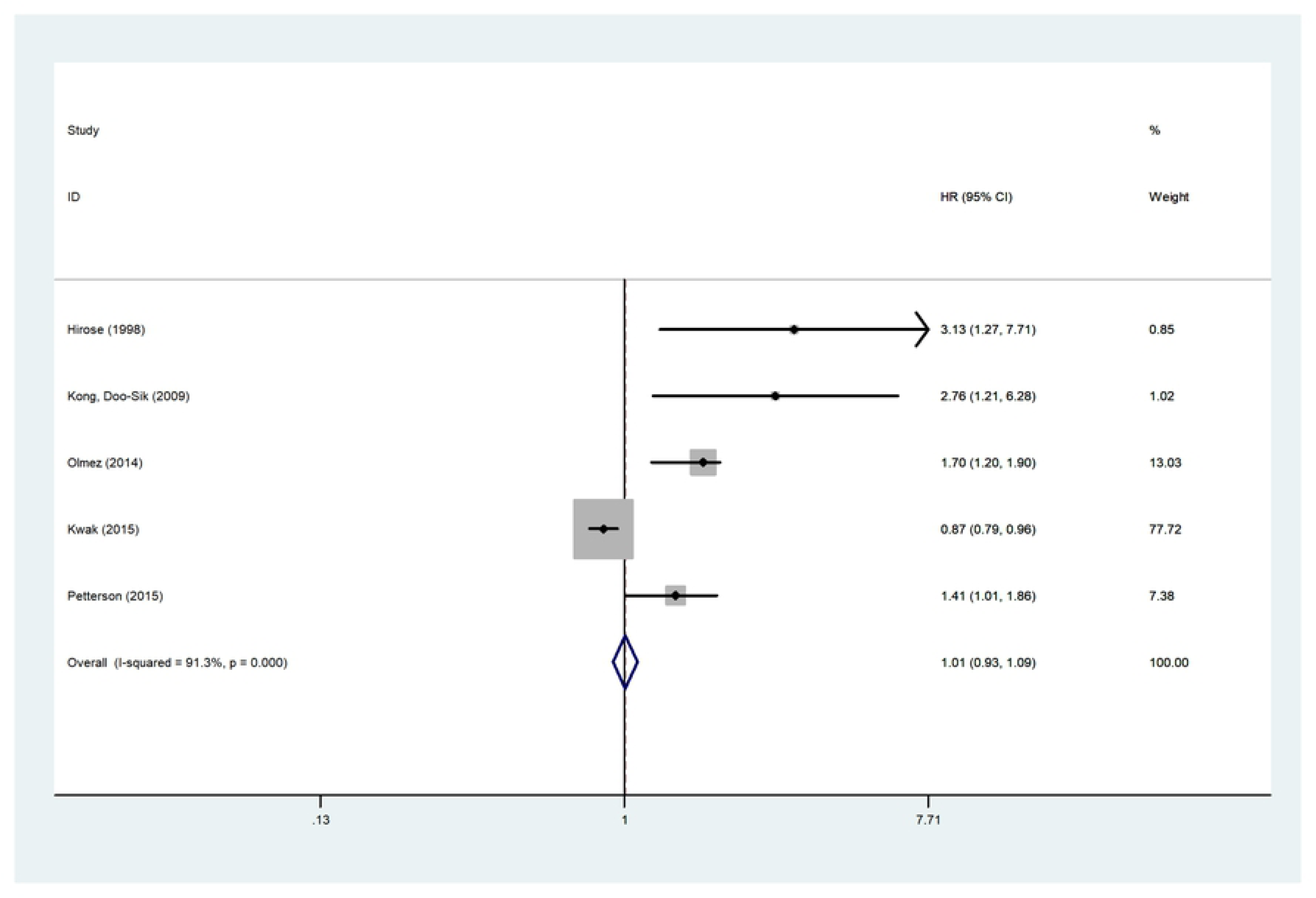
Forest plot of OS. The overexpression of c-Met was not associated with overall survival (HR=1.01, 95% CI:0.93-1.09)

#### Progression Free Survival

Kong 2009, Liu 2011, Olmez 2014 revealed the link between c-Met expression and progression free survival of glioma patients. In this meta-analysis, the pooled results indicated that the overexpression of c-Met was significantly associated with poor PFS (HR =1.92, 95% CI:1.42-2.58). There is low heterogeneity in this result (I^2^=28.3%, P=0.248). **(Figure 6)**

**Figure 6.**
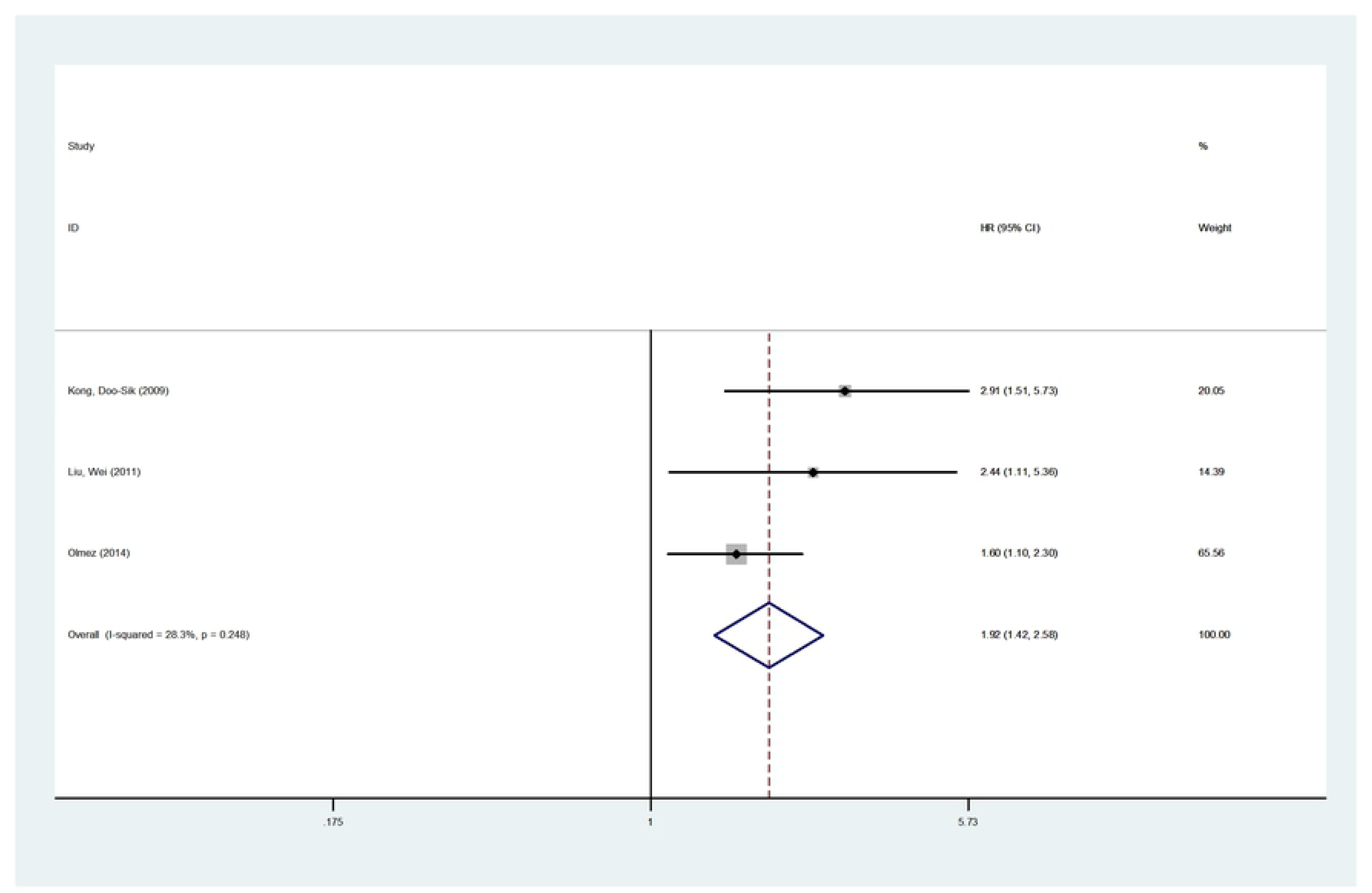
Forest plot of PFS. The overexpression of c-Met was significantly associated with poor PFS (HR=1.92, 95% CI:1.42-2.58).

#### Sensitivity analysis

To find the source of heterogeneity, sensitivity analysis was performed to validate the reliability of the conclusion, the results indicated that Kwak 2015 affect the overall HR for OS dramatically. This result indicates that if excluding Kwak 2015, the heterogeneity may decrease. **(Figure 7)**

**Figure 7.**
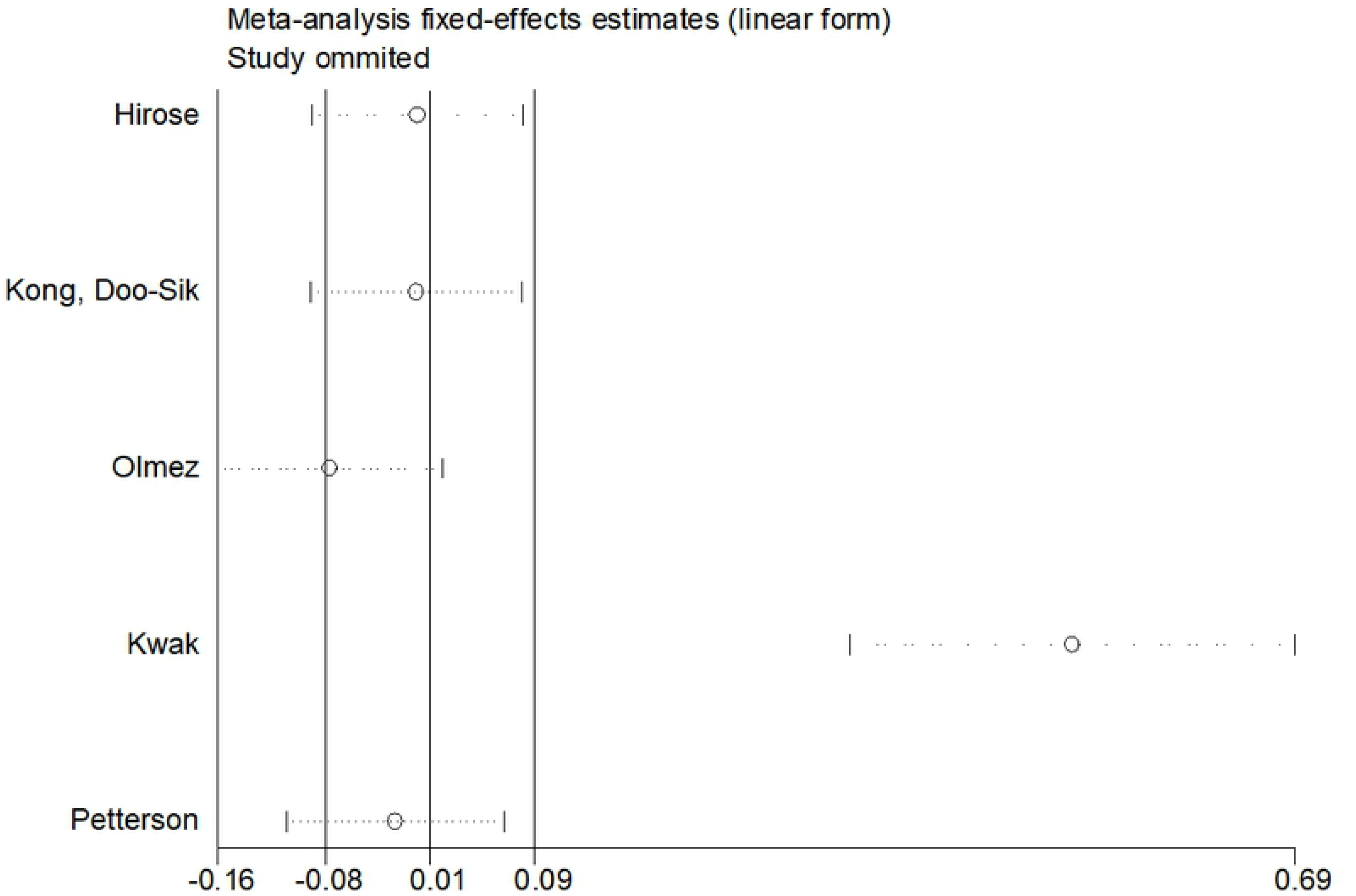
Sensitivity analysis of included studies. The results indicated that Kwak2015 affect the overall HR for OS dramatically.

#### Subgroup analysis

According to the areas mentioned in “Patients and Methods” of each article, we split data into two groups roughly. And then the subgroup analysis based on races was also performed. The results still showed manifest heterogeneity, the I^2^ (86.6% in Mongoloid, 0% in Caucasoid) just slightly decreased. **(Figure 8)**

**Figure 8.**
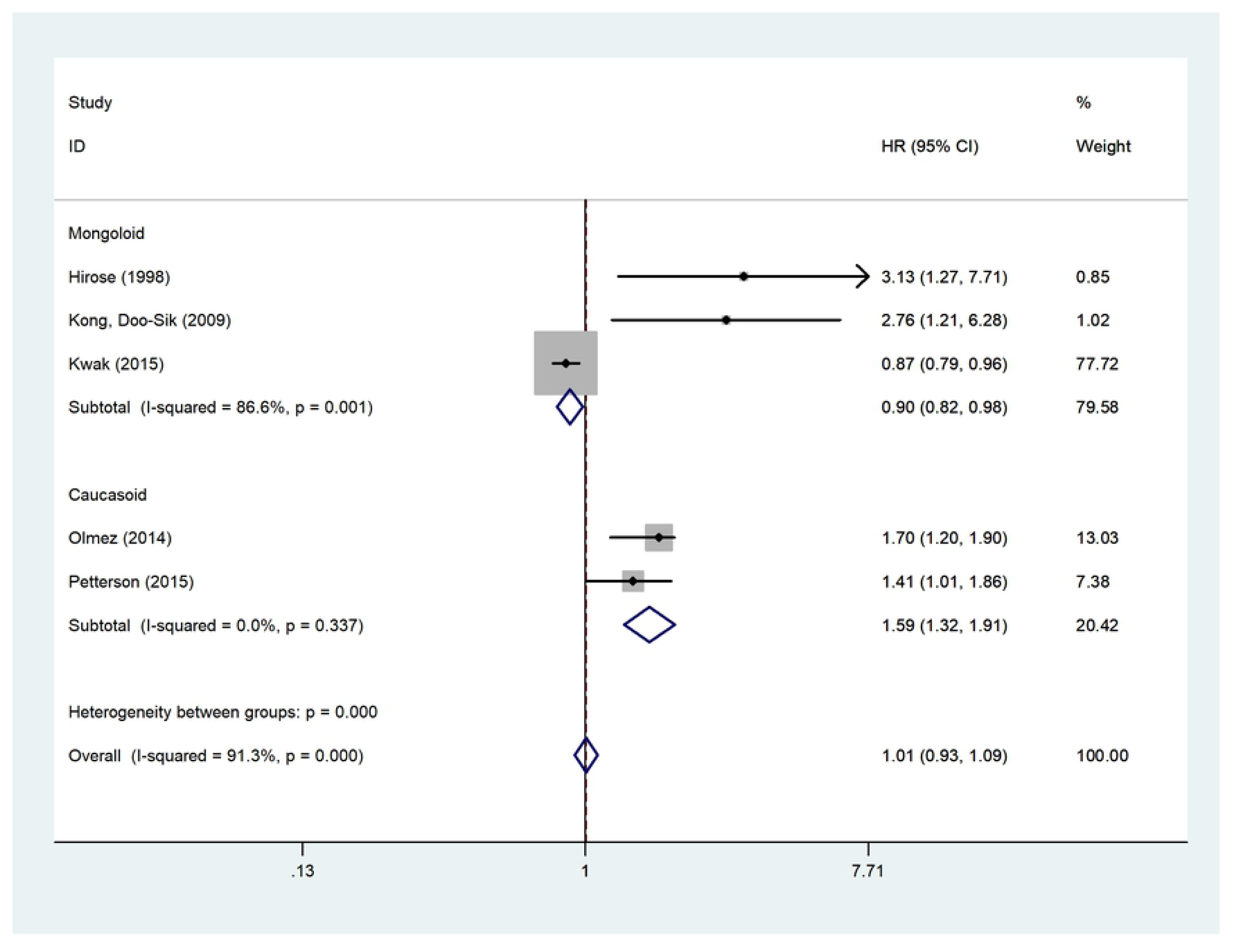
Forest plot of OS in patients races subgroup analysis.

#### Publication bias

Publication bias was analyzed by Begg’s Test with a P value of 0.462(P>0.05), by Egger’s Test with a P value of 0.056 (P>0.05), which suggested low publication bias among the included studies. **(Figure 9) (Figure 10)**

**Figure 9.**
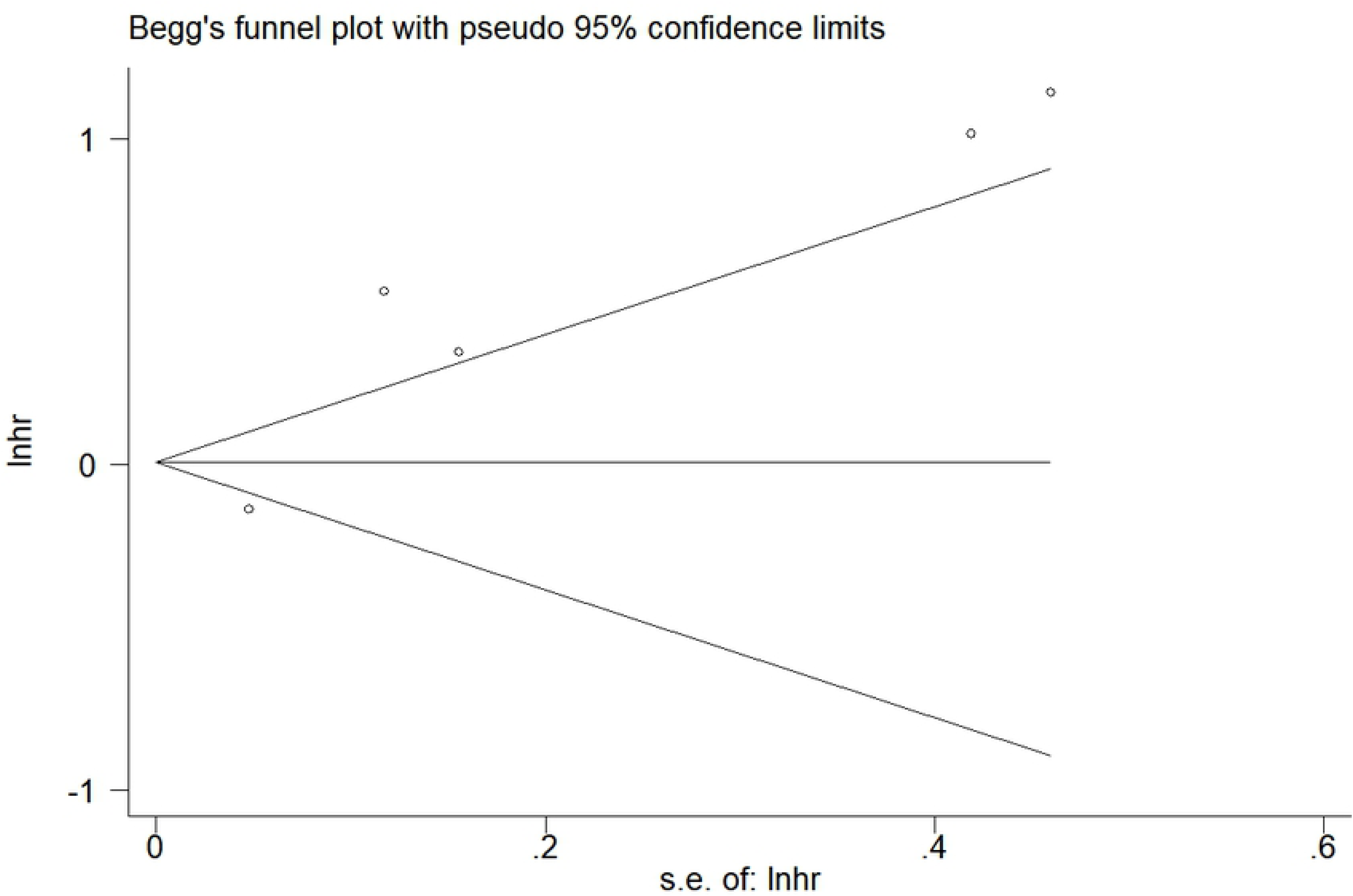
Publication bias of included studies. Begg’s Test. The results showed that there was low publication bias among studies.

**Figure 10.**
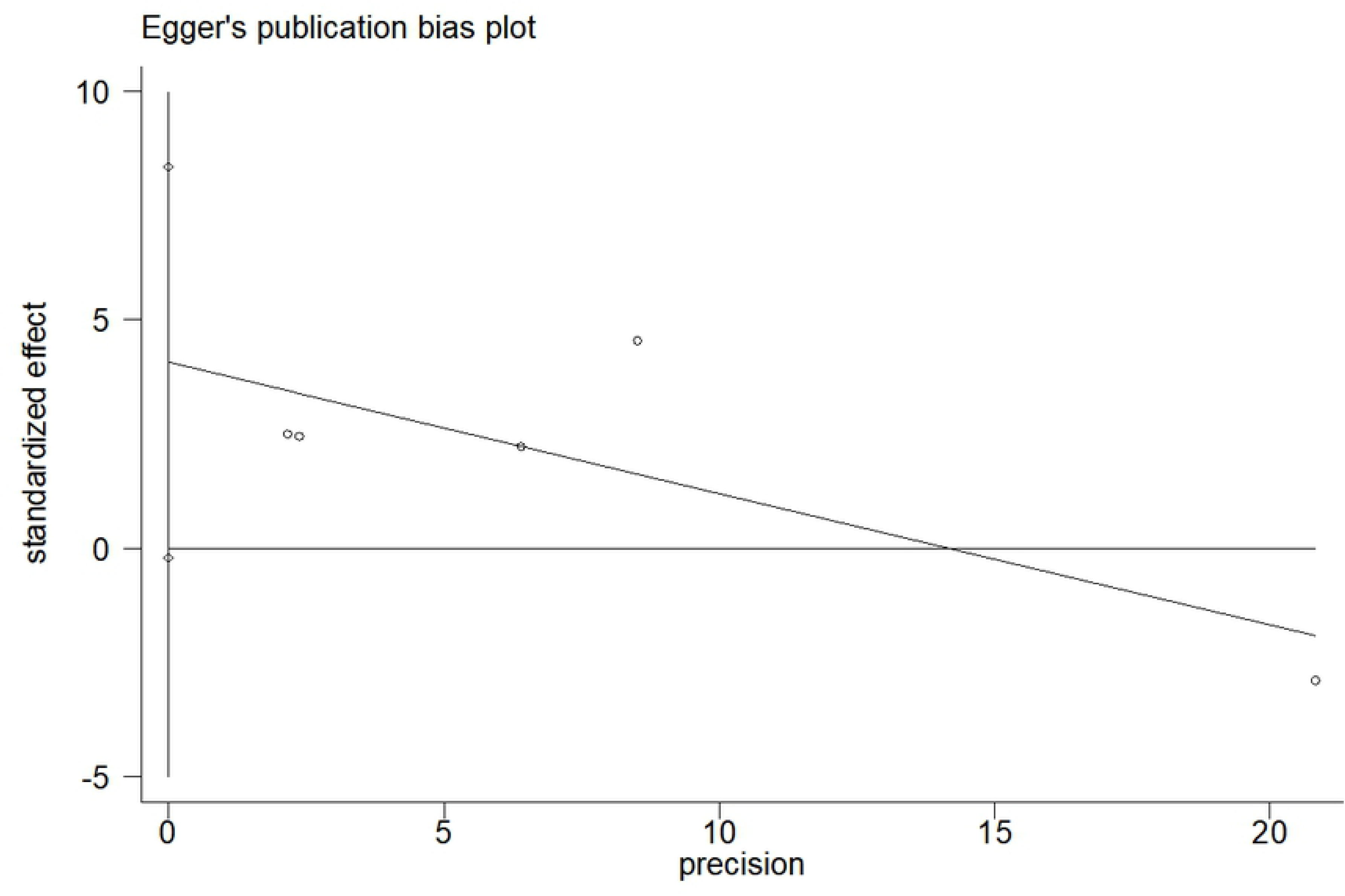
Publication bias of included studies. Egger’s Test. The results showed that there was low publication bias among studies.

## Discussion

High grade glioma is connected with its high mortality ^2^. Over the last few decades, despite significant advancements in diagnosis and multiple therapeutic approaches, high grade glioma is still known as its devastating pathology characteristics leading to the high recurrence rates and short overall survivals. With the rising of molecular biology and gene engineering, combinatorial treatments which target different molecular pathways and mechanisms is expected to improve the way we cure and diagnose this horrible disease ^24^. The HGF/MET signaling pathway is one of expected signal ways ^25, 26^. c-Met and its only ligand, hepatocyte growth factor (HGF), have been proved to be connected with many physiological and pathological processes ^27^. Alterations in HGF or c-Met are observed to be connected with more aggressive clinical behavior, contribute to cancer progression and influence patients’ prognosis in high grade glioma ^9, 28^. Therefore, finding out the relationship of c-Met and high grade glioma will contribute to facilitating the discovery of more effective diagnostic method, therapeutics and predicting prognosis.

The MET proto-oncogene, located on chromosome 7q21-31, encoded c-Met protein. In our study, Oncomine database analysis was firstly conducted to compare the MET copy number in glioblastoma and normal samples. These results supported that there indeed has difference between the MET copy number in glioblastoma patient samples and normal samples. Further analysis was conducted based on transcriptome data of CGGA. The results revealed that the high expression of c-Met was associated with the poor prognosis (OS and PFS) in glioblastoma patients (P<0.001),

Although in some original articles, Hirose 1998, Kong 2009, Olmes 2014, Petterson 2015 have observed significant relationship between overexpression of c-Met and poor prognosis of high grade glioma, but it is not convincible for the lack of samples. Also, there still existed some opposite views, like Kwak 2015, it holds the view that c-Met overexpression patients survived longer than patients without c-Met overexpression. So there are lots of controversies regarding to this problem. This situation prompted us to further explore the mechanism and underlying relationship between them.

Compared to small sample research, meta-analysis raises the accuracy and the credibility of the study as well as the conclusion. In order to further probe into this problem, we took this statistical method. Our meta-analysis has enrolled 6 studies, 503 patients, to investigate this problem. We finally arrived at a conclusion: the overexpression of c-Met was not strongly associated with overall survival (HR=1.01, 95% CI: 0.93-1.09). However, we noticed that Kwak 2015 had high weight in forest plot, which coincided with the results of sensitivity analysis, and deviated so broadly from other research in sensitivity analysis. In this case, to further probe into the problem, a meta-analysis excluding Kwak 2015 was conducted. The refined result and figure was shown in **Appendix 3**. The pooled result (HR=1.67, 95% CI:1.40-1.99) support the view that overexpression of c-Met has correlation with poor prognosis of high grade glioma patients. Meanwhile, the heterogeneity decreased obviously, the value of I^2^ declined to 33.4% from 91.3%, which validates that Kwak 2015 is one of the main causes of heterogeneity. In fact, Kwak 2015 may lack of experimental rigor, in its experiment, the number of experimental group (overexpression of c-Met) is 18, and the control group (not overexpression of c-Met) is 119. This big difference in two groups’ number may increase unnecessary contingency, which cause some potential errors in the test results.

As to detection method of c-Met, all articles have taken immunohistochemistry. Some articles had taken different cut-off value to define c-Met overexpression in this problem. Some articles, like Hirose 1998, Kong, Doo-Sik 2009, Liu 2011, Olmez 2014 took 30% as cut-off value. They defined <30% as negative, >30% as positive. Meanwhile, the latter three divided expression of c-Met into five grades: grade 0 – grade 5. Grade 0-1 (no expression, <30% expression) is negative. But for Petterson 2015, this article has experimented with different cut-off values. One cut-off was invalid while the other is effective. Meanwhile, Petterson 2015 has held that c-Met overexpression is a time-dependent factor to glioblastomas prognosis, which could only be observed in survived more than 8.5 months’ patients. However, other articles didn’t mention this information. To our disappointment, because we don’t have enough data, we can’t do more analysis about these problems. More detailed researches are expected to explore the exact role that time factor play in this problem.

We also use forest plot to ensure the relationship between c-Met overexpression and progression free survival of glioma patients. Kong 2009, Liu 2011, Olmez 2014 was enrolled in the study. The results showed that the overexpression of c-Met was significantly associated with poor PFS (HR =1.92, 95% CI:1.42-2.58).

In the following, we tried to find out the sources of heterogeneity by performing subgroup analysis according to races. In the subgroup analysis based on race classification, the heterogeneity just fell down a bit. In Caucasoid group, heterogeneity fell down dramatically (I^2^=0). In Mongoloid group, the heterogeneity was still high. However, when Krak 2015 was excluded, the result shows low heterogeneity in two groups (0% in Mongoloid, p=0.84; 0% in Caucasoid, p=0.337). The figure was shown in **Appendix 3**. The high heterogeneity in direct sequencing is mainly caused by Kwak 2015, which deviated from the other studies with high HR and weight. Because of the lack of detailed ethnicity data, this subgroup was simply based on the address of hospital mentioned in the “materials and methods” part of each article, but not the exact race of each patient, which naturally resulted in biased conclusion. It will be too hasty to simply affirm or negate the role of this factor played in final conclusion, more detailed subgroup research should be conducted in the future.

Moreover, we have also analyzed publication bias of enrolled studies. According to Begg’s test and Egger’ test, there was low publication bias among the included studies (p>0.05). This made conclusion more incredible.

Our study shows strength as follows: Firstly, this article is the first meta-analysis that investigates the connection between c-Met overexpression and high grade glioma patients survival. Second, this article has performed this diversification analysis, including Oncomine, transcriptome and meta-analysis to strengthen the credibility of conclusion.

There were still several limitations in our work. Because of the lack of original information, we can’t do more detailed subgroup analysis, like whether 8.5 month could influence the relationship between the expression of c-Met and the prognosis of high grade glioma patients. Significant statistical heterogeneity still existed in the current study. Therefore, in order to evaluate prognostic significance of c-Met more precisely, larger sample sized, multi-center and cohort studies should be performed in the future.

In summary, with the foundation of the above analysis, we demonstrated that compared with normal samples, MET expression level was expressed higher in glioblastoma patient samples. The MET proto-oncogene encoded c-Met protein, moreover, overexpression of c-Met indeed was correlated with poor prognosis of high grade glioma patients.

## Acknowledgement

This work was supported by the National Nature Science Foundation of China (Grant nos. 81571737) and the Project of the Jilin Provincial Science and Technology Department of China (Grant nos. 20130204028GX and 20140413037GH).

---

Table 1 Characteristics of the studies included in the meta-analysis.

Table 2 Details of the Oncomine database analysis

